# Transcranial Focused Ultrasound Neuromodulation of Voluntary Movement-related Cortical Activity in Humans

**DOI:** 10.1101/2020.05.18.103176

**Authors:** Kai Yu, Chang Liu, Xiaodan Niu, Bin He

## Abstract

Transcranial focused ultrasound (tFUS) is an emerging non-invasive brain stimulation tool for safely and reversibly modulating brain circuits. The effectiveness of tFUS on human brain has been demonstrated, but how tFUS influences the human voluntary motor processing in the brain remains unclear. We apply low-intensity tFUS to modulate the movement-related cortical potential (MRCP) originating from human subjects practicing a voluntary foot tapping task. 64-channel electroencephalograph (EEG) is recorded concurrently and further used to reconstruct the brain source activity specifically at the primary leg motor cortical area using the electrophysiological source imaging (ESI). The ESI illustrates the ultrasound modulated MRCP source dynamics with high spatiotemporal resolutions. The MRCP source is imaged and its source profile is further evaluated for assessing the tFUS neuromodulatory effects on the voluntary MRCP. Moreover, the effect of ultrasound pulse repetition frequency (UPRF) is further assessed in modulating the MRCP. ESI results show that tFUS significantly increases the MRCP source profile amplitude (MSPA) comparing to a sham ultrasound condition, and further, a high UPRF enhances the MSPA more than a low UPRF. This work provides the first evidence of tFUS enhancing the human voluntary movement-related cortical activity through excitatory modulation.

## 1. Introduction

As an emerging non-invasive neuromodulation tool, the low-intensity transcranial focused ultrasound (tFUS) delivers highly controllable mechanical energy from transducers, penetrating the skull with relatively low tissue attenuation and modulates the targeted brain circuits with high spatial selectivity. tFUS is featured with a vast parametric space, and by tuning its parameters, the ultrasound wave can be steered and be used to produce excitatory or inhibitory neural effects. Such neuromodulatory effects of tFUS for effective brain stimulation have been evidenced on a wide range of animal models (Dallapiazza et al., 2018; Daniels et al., 2018; Folloni et al., 2019b; Lee et al., 2016c; Mohammadjavadi et al., 2019; Tufail et al., 2011; Verhagen et al., 2019; Yang et al., 2018; Yoon et al., 2019; Yu et al., 2016) and, recently, demonstrated in humans (Lee et al., 2016a; Lee et al., 2015; Lee et al., 2016b; Lee et al., 2017; Legon et al., 2018a; Legon et al., 2018b; Legon et al., 2014). For instances, the low-intensity diagnostic ultrasound energy has been shown to suppress the pain, improve the subjects’ mood and global affect (Hameroff et al., 2013). A recent study on using ultrasonic thalamic stimulation assisted a patient to recover consciousness after severe brain injury (Monti et al., 2016).

Among accumulating experimental demonstrations on healthy humans, a pioneer study using low-frequency and low-intensity tFUS was able to achieve robust neuromodulation effects at the primary somatosensory cortex (S1), evidenced through the ultrasound-modulated sensory-evoked brain activity recorded at electroencephalogram (EEG) sensors and the enhancement of sensory discrimination abilities (Legon et al., 2014). Another pilot study on tFUS modulating the S1 circuits reported direct evoked limb sensations and the ultrasound stimulation event-related potentials (ERPs) specifically at C3 and P3 EEG electrodes (Lee et al., 2015). Later, the acoustic stimulation was ipsilaterally targeted at S1 and S2 (i.e. the secondary somatosensory cortex) simultaneously by multiple transducers. Extensive tactile sensations were elicited and reported by the human subjects (Lee et al., 2016a).

The visual cortex is also a well-studied brain target for tFUS research. On human, researchers administered sonication to the primary visual cortex (V1). From the concurrent blood-oxygenation-level-dependent (BOLD) functional magnetic resonance imaging (fMRI), researchers observed ultrasound-elicited brain activations at the targeted V1 as well as the related visual and cognitive brain network. Sonication-mediated ERPs and phosphene perception were also reported (Lee et al., 2016b). Very recently, the ultrasound-induced illusory visual percepts were further investigated by applying repeated transcranial diagnostic ultrasound at visual cortical region identified by transcranial magnetic stimulation (TMS) eliciting phosphene (Schimek et al., 2020).

The motor cortex has been one of the popular sonication targets on animal models for eliciting direct brain activation by tFUS, demonstrated by more direct motor response readout (Kim et al., 2014; King et al., 2013; Mehic et al., 2014; Tufail et al., 2010; Ye et al., 2016; Yuan et al., 2020) and/or measurements from the corresponding muscle activities using electromyogram (EMG) (King et al., 2013; King et al., 2014; Qiu et al., 2017; Yoon et al., 2019). In spite of this, the demonstrations of tFUS modulation on human motor cortex are somehow limited. One of the recent efforts was to employ a 7-Tesla BOLD fMRI for observation of tFUS modulation in the targeted finger representation areas in terms of neurovascular signal change while the subjects performed a simultaneous cued finger tapping task. The activation volume of the thumb motor representation area was significantly increased by the tFUS neuromodulation (Ai et al., 2018). Another effort was to further test how the ultrasound effects the motor cortical excitability and behavior output by developing a novel hybrid brain stimulation device for concurrent transcranial ultrasound and magnetic stimulation (Legon et al., 2018b). The tFUS neural inhibitory effects were inspected and quantified through the amplitude reduction of single-pulse motor-evoked potentials and the attenuation of intracortical facilitation. Interestingly, the tFUS significantly shortened the reaction time on a stimulus response task (Legon et al., 2018b).

However, our current understanding on how tFUS can influence the human voluntary motor processing in the brain is still lacking. In this study, we test the ultrasound neuromodulatory effects on movement-related cortical potential (MRCP) which occurs during the preparation for and execution of movement with clinical significance on rehabilitation in patients (Niazi et al., 2012; Shakeel et al., 2015; Spring et al., 2016; Xu et al., 2014), scientific research on studying motor skill learning (Wright et al., 2011), and enhancing the brain-computer interface (BCI) (Dremstrup et al., 2013; Liu et al., 2020). Beyond the EEG delineation at the sensor level, in this study, we employ scalp EEG based electrophysiological source imaging (ESI) (He and Ding, 2013; He et al., 2018; Yu et al., 2016) to model the MRCP at the corresponding motor cortical region in healthy humans, and further evaluate the MRCP change at the EEG source domain for an enhanced spatial specificity.

Furthermore, by virtue of computer modeling and simulations (Plaksin et al., 2016), the ultrasound pulse repetition frequency (UPRF) has been deemed as one of the critical ultrasound parameters to achieve excitatory/inhibitory neural effects. In a recent study, the acoustic radiation force produced by the UPRF has been inferred as the most crucial source of behavioral responses (Kubanek et al., 2018). Most recently, the UPRF was also found as a critical parameter in balancing the excitatory/inhibitory neuronal activities (Yu et al., 2019). In this study, we further assess the role of UPRF in modulating the human MRCP.

## 2. Materials and Methods

### Participants

Fifteen healthy human subjects were recruited in this experiment (10 males, 5 females, mean age of participants: 33.39 ± 14.02 years). Each subject attended on-site interview for safety screening and provided written informed consent before participating the experiment. The study was reviewed and approved by the Institutional Review Board at Carnegie Mellon University.

### Experimental Setup

Prior to the tFUS-EEG session, each participant received a 3-Tesla magnetization-prepared rapid gradient-echo (MP-RAGE) T1-weighted structural magnetic resonance imaging (MRI, Siemens Verio) in order to establish high-resolution individual brain anatomical models. The models were later used to identfy the brain target through FreeSurfer cortical reconstruction (Dale et al., 1999; Desikan et al., 2006) for guidance of the low-intensity focused ultrasound focus.

During the tFUS-EEG session, the subject was seated inside a sound and electromagnetic shielding room (IAC Acoustics, USA) and was instructed to wear foam earplugs (3M, USA) during experimental sessions. A 24-inch LCD monitor with a viewing distance of 50 cm was used to instruct each subject to start or stop voluntary foot pedaling (illustrated in Figure 1). During the task, an accelerometer (ADXL335, Adafruit Industries LLC, USA) mounted on a foot pedal (Foot Control 704NS-GR, Elna Machine Parts, USA) detected and transmitted the fast motion of foot pressing to ExG AUX Box (Brain Products GmbH, Germany). Concurrent 64-channel EEG data were acquired using BrainAmp (Brain Products GmbH, Germany), with electrode positions FCz and AFz chosen as reference and ground. The electrical impedance of all electrodes was kept below 10 kΩ during the EEG capping. Positions of electrodes were digitized over each subjects’ scalp using an optical-based EEG PinPiont system (Localite GmbH, Germany). The ultrasound transducer was held and mounted on top of the EEG cap using a 3D-printed helmet. An optical-based brain navigation system (Localite GmbH, Germany) was utilized with the input of the structural MRI data and an optical marker attached over the forehead to track and guide the position and orientation of the ultrasound transducer in real time. Surface adhesive electrodes (Medi-trace 530 series, Covidien, USA) were applied onto the skin of calf and were connected to ExG AUX Box (Brain Products GmbH, Germany) for monitoring muscle activities at the lower leg for validating each foot pedaling motion.

**Figure 1.**
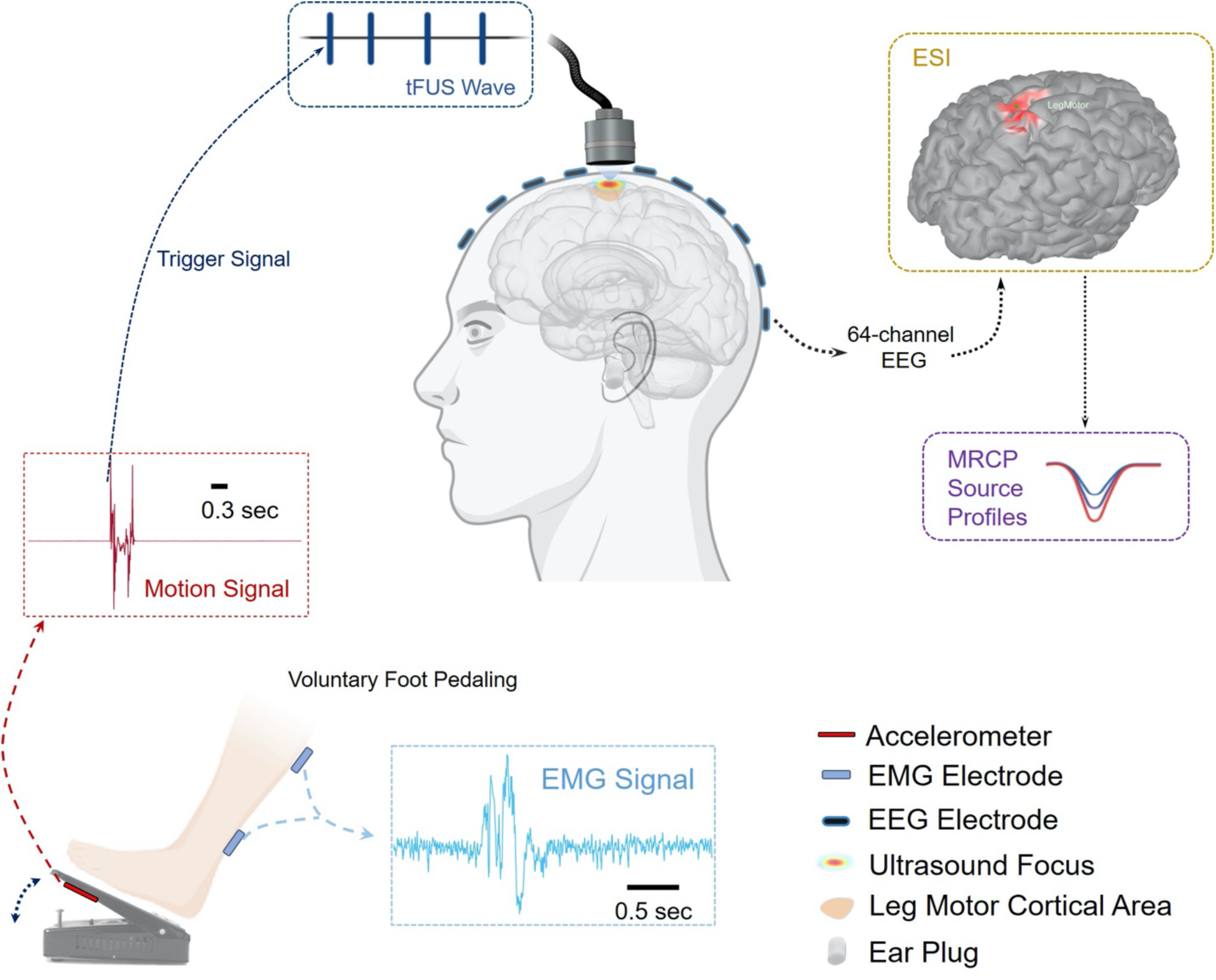
Diagram of the overall experimental setup. The voluntary right foot pedaling generates the motion signal, and only the onset of this motion signal (from a fast-response, high-precision comparator) is detected as the trigger signal for transmitting transcranial focused ultrasound (tFUS) onto the primary leg motor area. The concurrent 64-channel electroencephalogram (EEG) is recorded for electrophysiological source imaging (ESI). The reconstructed source activity at the leg motor cortical area is further extracted and the movement-related cortical potential (MRCP) source profiles are thus derived for further assessing the tFUS neuromodulatory effects. The electromyogram (EMG) is recorded simultaneously for monitoring and validating successful voluntary foot pedaling trials. The subject is instructed to wear earplugs throughout each testing session.

### Experiment Procedures

The overall experimental design is shown in Figure 1. The motion task was repeated in three sessions with respective ultrasound setup in a random order (see details in *Ultrasound Setup*). The subjects were instructed to initiate pressing the foot pedal using their right leg roughly every 3-5 seconds on their own discretion after the screen turned to green color. About 120 trials of right foot pedaling were conducted and recorded. The subjects were instructed to keep the upper body still and eyes open. The subjects stopped foot motion once the screen turned to red.

### Ultrasound Setup

A single-element focused ultrasound transducer with an acoustic aperture diameter of 25.4 mm and a nominal focal distance of 38 mm (AT31529, Blatek Industries, Inc., USA) was used in this study. A 3D-printed collimator (made in VeroClear resin) was attached to the transducer in order to match the focal length of the transducer with the estimated physical distance from the acoustic aperture to the targeted motor cortex. The ultrasound signal was generated by two function generators (33220A, Keysight Technologies, Inc., USA) and amplified by a radiofrequency (RF) power amplifier (2100L, Electronics & Innovation, Ltd., USA), driving the ultrasound transducer. The first function generator was triggered by an output signal from a home-made circuit based on a fast and precise voltage comparator (AD790, Analog Devices Inc., USA), which was monitoring the output from the accelerometer in real time (shown in Figure 1). This trigger signal was further stretched to 5 msec long by TriggerBox (Brain Products GmbH, Germany) for synchronization with EEG recordings. Once triggered, the first function generator was used to create the UPRF and determine the number of pulses. The second function generator was triggered by the output of the first one in burst mode and generated ultrasound fundamental frequency (UFF) of the sinusoidal waveforms and determined the number of cycles per pulse (CPP).

As illustrated in Figure 2A, this study used a UFF of 0.5 MHz, CPP number of 100. Each sonication lasted for 500 msec with two levels of UPRF, i.e. 300 and 3000 Hz practiced in two sessions (denoted as “UPRF 300Hz” and “UPRF 3000Hz”, respectively). The ultrasound spatial peak pressure applied to the scalp was measured as 809.2 kPa (shown in Figure 2B-C, spatial-peak pulse-average intensity I_SPPA_: 5.90 W/cm^2^), with an estimated ultrasound pressure of 288.3 kPa (Figure 2D, I_SPPA_: 1.17 W/cm^2^) arriving at the targeted cortical brain with an acoustic insertion loss of −8.96 dB. This pressure estimation was based on a 3-dimensional *ex-vivo* transcranial ultrasound scanning using a needle hydrophone (HNR-0500, Onda Corp. USA) placed in a full real human skull sample (OK-14472, Skulls Unlimited International, Inc., USA) and driven by a 3-axis precision motion system (BiSlide system, Velmex Inc., USA). From Figure 2B, it can be seen that the −3dB contour of the focus is about 3 mm, and this focal size makes the administered tFUS eligible to spatially targeting at the leg motor cortical area. A sham tFUS session was also introduced for rigorous comparisons. In this sham condition (denoted as “Sham US”), the effective acoustic aperture was physically detached from subject’s scalp for 4-6 centimeters while the transducer was still transmitting ultrasound at the UPRF of 3000 Hz.

**Figure 2.**
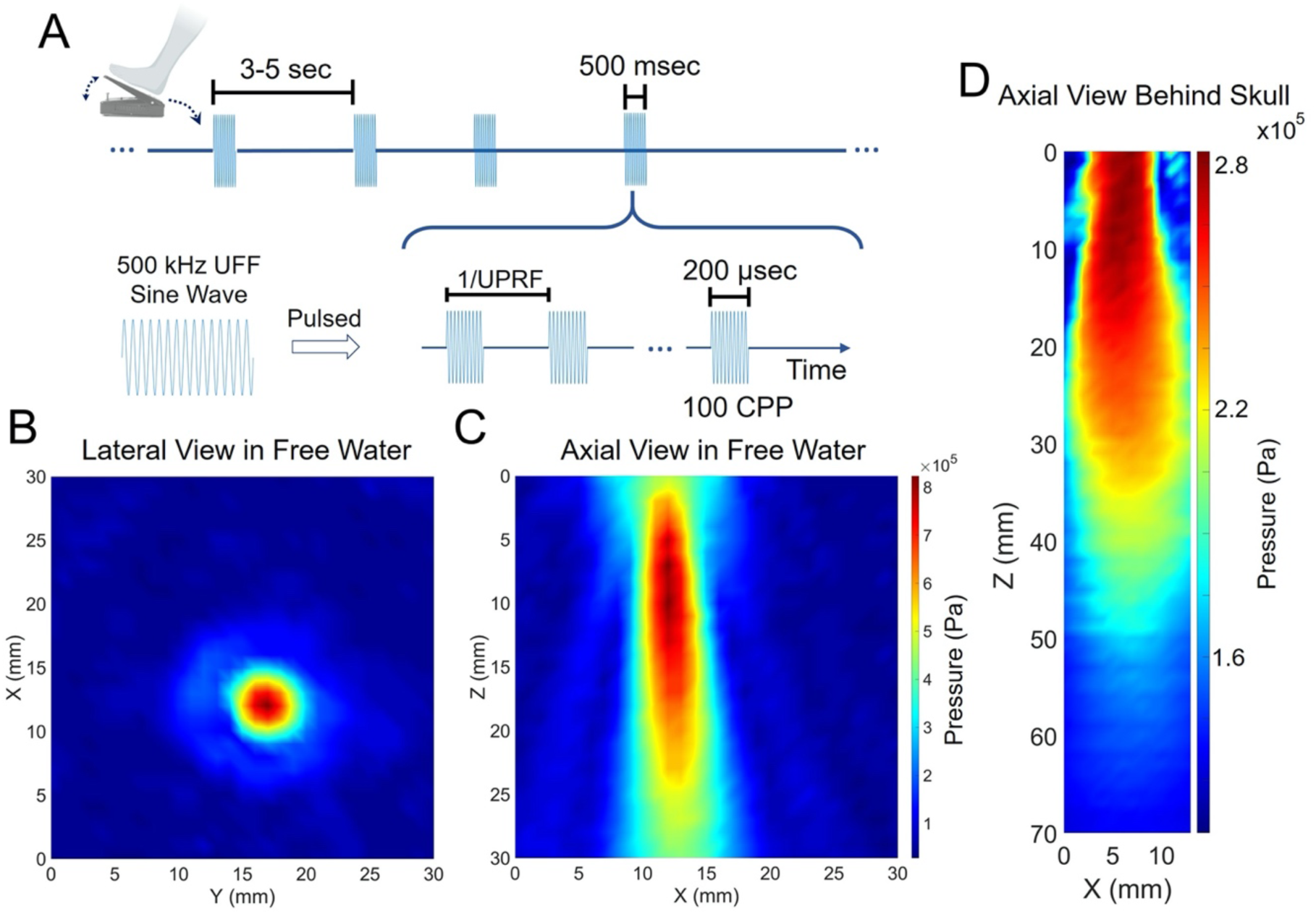
Temporal and spatial profiles of the ultrasound for modulating the MRCP. (A) The temporal waveforms of the pulsed tFUS triggered by the foot pedaling motion. (B-C) The lateral and axial views of the ultrasound focus and beam in free water measured with a 3D pressure scanning system. (D) The axial view of the transcranial ultrasound beam within a full real human skull sample. The smaller Z number along the axial view represents the shorter distance from the transducer’s acoustic exit plane.

### Electrophysiological Signal Processing

The lower leg EMG, sampled at 5 kHz, was used to assess the quality of each foot pedaling trial. During the offline data processing, we scrutinized the signals from the EMG and the accelerometer demonstrated in Figure 1, and removed the trials reported from the accelerometer while the EMG signals of those trials were not validated as a normal foot pressing activity. This step was to exclude unintentional foot movement, like foot slipping or uncompleted foot pedaling.

The EEG was sampled at 5 kHz and filtered using a bandpass filter with the lower cut-off frequency at 1 Hz and the higher cut-off frequency at 45 Hz. The pre-stimulus period was set as 400 msec before the trigger signal, and the period of 600 msec after the onset of the trigger signal was deemed as post-stimulus period in EEG individual epoch. Independent component analysis (ICA) (Makeig et al., 2002) and/or signal-space projection (SSP) (Uusitalo and Ilmoniemi, 1997) were used to identify and clean artifacts, mainly the strong eye blinking during the voluntary movement. The MRCPs in the time domain were normalized against the first 100 msec during the pre-stimulus period. The 1-second EEG epochs were then averaged across the trials for each experimental condition by aligning the detected trigger signal (i.e. Time 0 in Figure 3A, D, G, Figure 4A and Figure 6). For EEG-based source modeling and imaging, the boundary element method (BEM) (Hämäläinen and Sarvas, 1989; He et al., 1987) head model for each human subject was established using OpenMEEG (Gramfort et al., 2010), which consisted of three layers, i.e. scalp, skull and brain with relative conductivities of 1, 0.0125 and 1, respectively. The minimum norm imaging (MNI) was used to solve the inverse problem, thus reconstructing the cortical source activity. The source activity was imaged using the MNI for the duration of sonication. Further, we took a source patch with an area of 3.2 – 3.5 cm^2^ from the primary leg motor area of reconstructed MRCP source activity. The measurement of MRCP source was the averaged activity across the patch. The temporal dynamics of the MRCP measurements were depicted as a time profile within the 1-sec epoch period. The peak-to-peak amplitude of MRCP source profile was then used for comparisons and statistical analyses. The EEG signal processing was performed with Brainstorm toolbox (Tadel et al., 2011) in MATLAB R2018a (Mathworks, Inc., USA).

**Figure 3.**
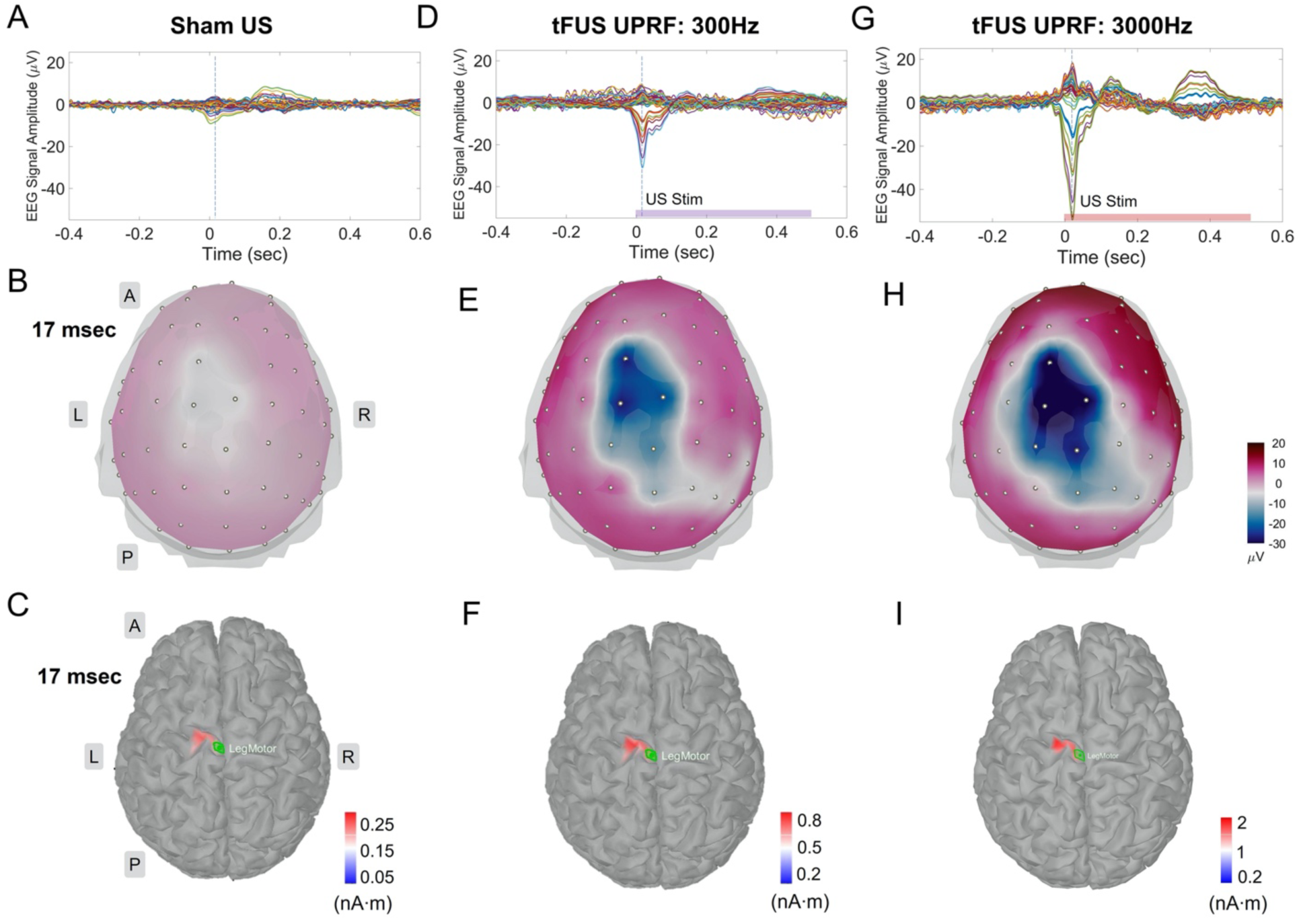
The MRCP source analyses of the 64-channel scalp EEG acquired from Sham US (A), UPRF 300 Hz (D) and UPRF 3000 Hz (G) sessions. The vertical dashed lines in the EEG butterfly plots indicate the time point, i.e. 17 msec. The 3D EEG voltage topography maps (top view) show a whole-brain comparison between the three ultrasound conditions using the same color scale (B, E and H). The panels of (C), (F) and (I) show more detailed views of the brain activities specifically at the primary motor cortex (M1) in responses to the three ultrasound sessions, respectively. The ESI illustrated reconstructed current source densities. The green patches in these ESI panels represent the seed areas for further extracting the MRCP source profiles. Different color scale bars are used to allow better presentation of the localized motor source at the cortical brain. L: left, R: right, A: anterior, P: posterior.

**Figure 4.**
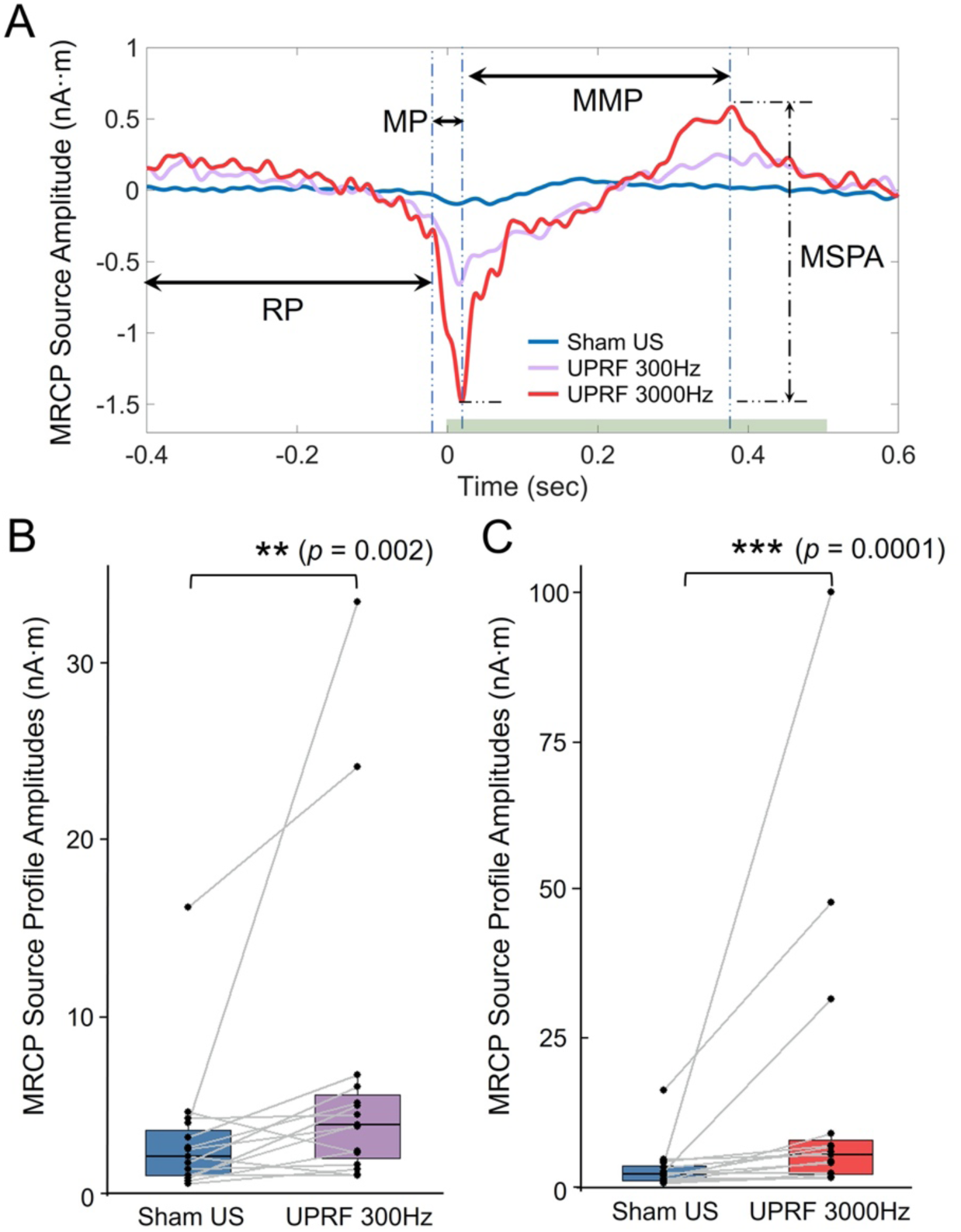
The tFUS modulates the MRCP by strengthening the MRCP source profile amplitudes (MSPA). (A) The MRCP source profiles (MSP) obtained from the Sham US, UPRF 300Hz and UPRF 3000Hz sessions are illustrated. The typical three MRCP components are indicated with horizontal arrows. RP: readiness potential. MP: motor potential. MMP: movement-monitoring potential. The MSPA is measured from the MSP of each conditions. The horizontal green bar represents the sonication duration. (B-C) Data are shown in the boxplots as the median with 25% and 75% quantiles (lower and upper hinges). Statistics by one-tail non-parametric paired Wilcoxon rank sum test for examining the effect of the tFUS with UPRF of 300 Hz (B) and the tFUS with UPRF of 3000 Hz (C) on the MSPA. \*\**p* < 0.01, \*\*\**p* < 0.001.

**Figure 5.**
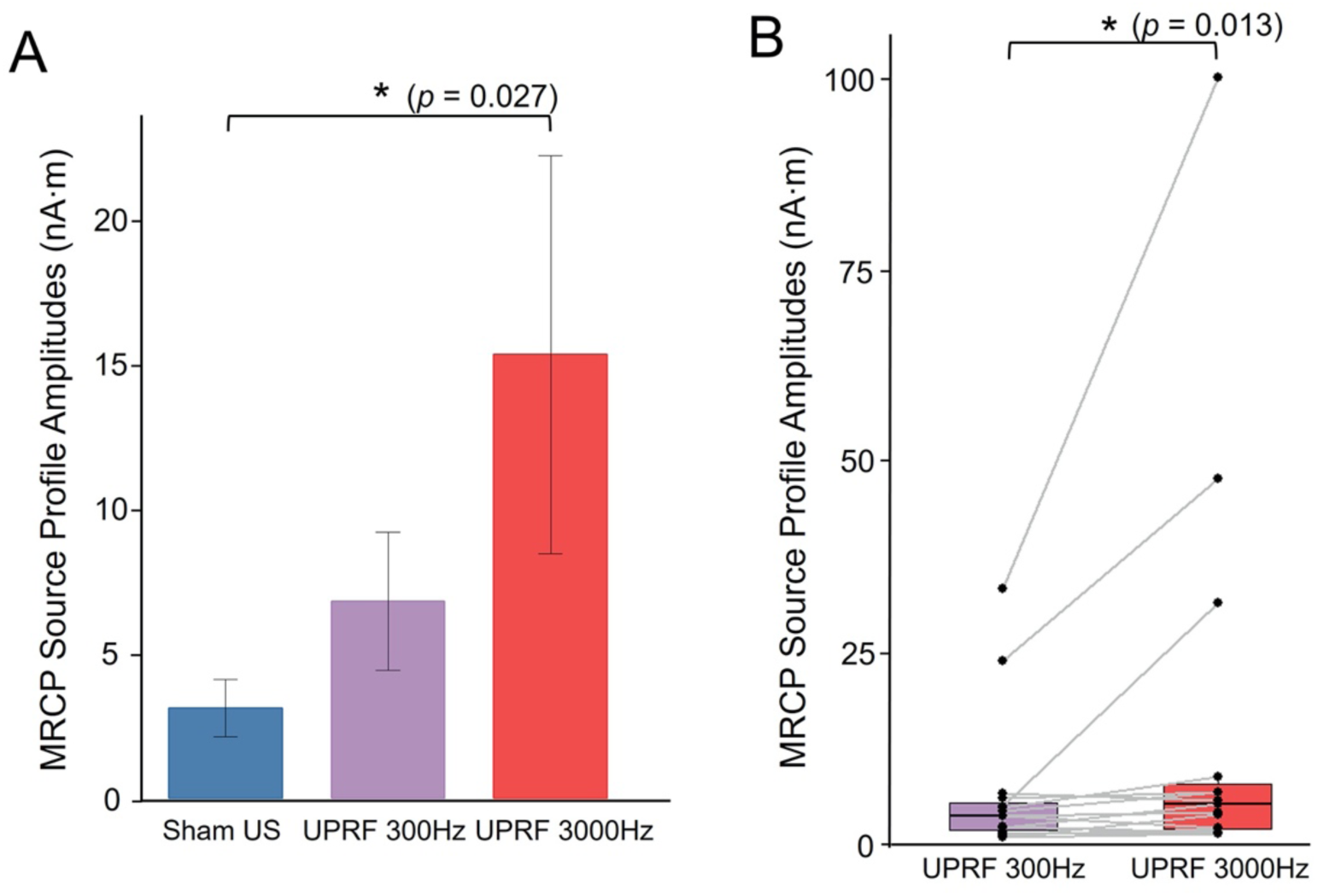
Examining the neural effects of tFUS UPRF. (A) Data are shown as the mean±s.e.m., statistics by Kruskal-Wallis rank sum test. \**p* < 0.05. (B) Data are shown in the boxplots as the median with 25% and 75% quantiles (lower and upper hinges). Statistics by one-tail non-parametric paired Wilcoxon rank sum test for examining the effect UPRF increase. \**p* < 0.05.

**Figure 6.**
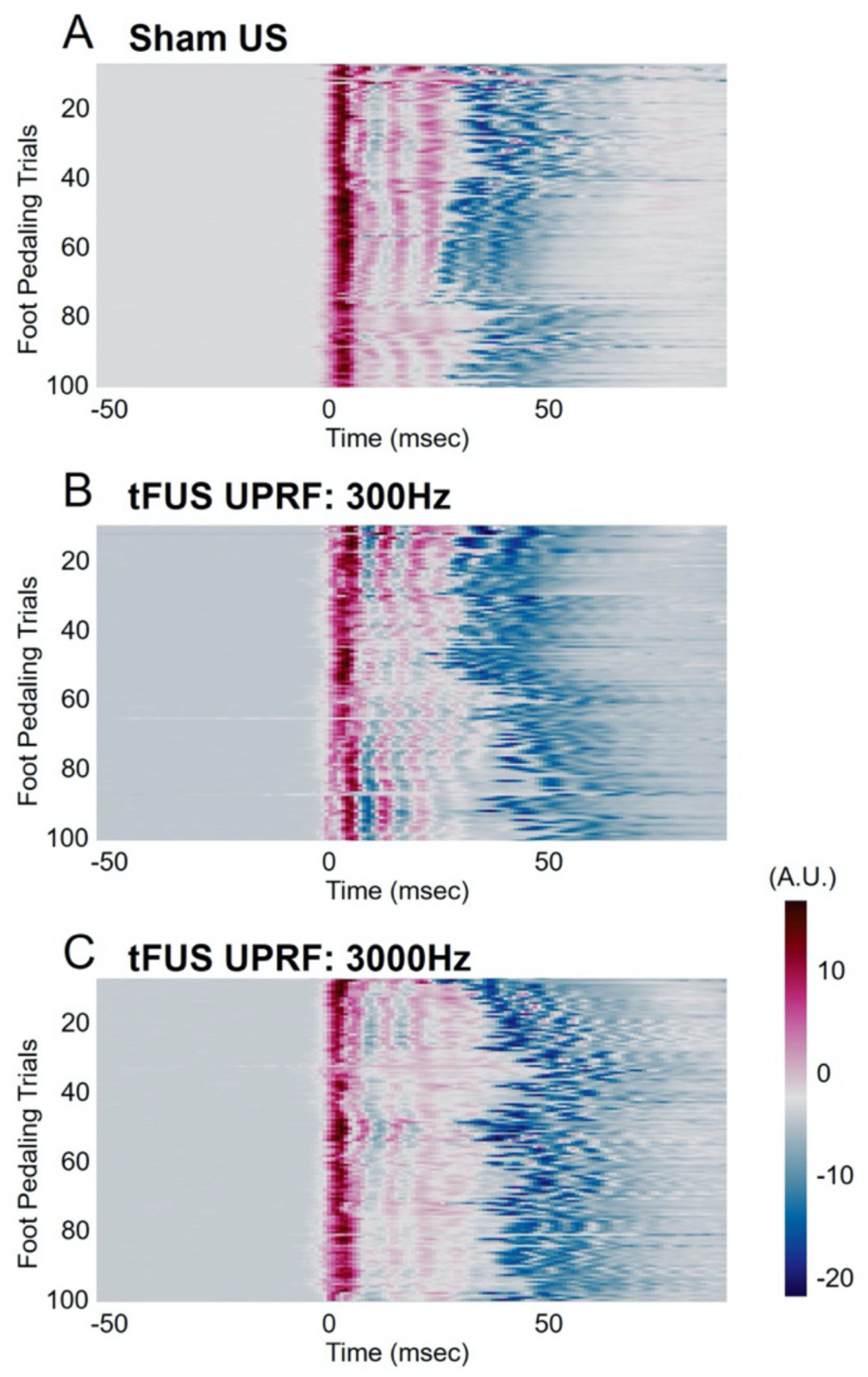
The comparisons of foot pedaling motion signals across all the trials in each ultrasound session, i.e. Sham US (A), UPRF 300Hz (B) and UPRF 3000 Hz (C). Arbitrary units (A.U.) are used for describing the motion signals.

### Statistical Analyses

Our statistical analyses focused on the tFUS modulation of the amplitude of MRCP source profiles. The first null hypothesis to be tested is that the MRCP source profile has no greater amplitude in the tFUS conditions than that in the sham condition. For testing this hypothesis, we performed one-tail non-parametric paired Wilcoxon rank sum test to examine the tFUS effects. The second null hypothesis is that the UPRF change will have no effect on the MRCP source amplitude. The Kruskal-Wallis rank sum test was performed to examine this hypothesis. Next, paired Wilcoxon test was further employed to assess whether the increased UPRF will lead to a stronger neuromodulation effect. For both statistical tests above, we performed Shapiro-Wilk test for examining data normality. All the above statistical tests were conducted in R V3.2.1. In addition, due to the correlation between the MRCP amplitude and movement strength, we investigated whether there were significant differences regarding the foot pedaling strength across different sessions for each subject, we did non-parametric permutation-based tests on the motion and EMG signals for ruling out such a possible confound. These additional statistical analyses were performed with Brainstorm toolbox (Tadel et al., 2011).

## 3. Results

The multi-channel EEG butterfly plots in Figures 3A, D and G illustrate a significant increasing in terms of sensor-level MRCP amplitude due to the presence of tFUS at the left primary leg motor area. Such EEG signal amplitude increases were specifically detected at electrodes C1, FC1, Cz, CPz and CP1, located close to the targeted brain area. In the Sham US condition only with right foot pedaling, the EEG voltage topography map reflected a mild activation at 17 msec majorly at the left brain hemisphere (Figure 3B).

By applying the EEG-based source imaging, we further localized the MRCP source activity at the precentral gyrus of left hemisphere, i.e. primary motor area. The reconstructed MRCP source covered a region with an approximate area of 4 cm^2^ and exhibited a current source density (CSD) amplitude of 0.27 nA·m. This MRCP-related CSD amplitude was increased to 0.8 nA·m after the tFUS (UPRF 300Hz) has been directed to the primary leg motor area (Figure 3F). By increasing the UPRF to 3000 Hz while maintaining the sonication duration, the MRCP was further enhanced at both the sensor level (Figure 3G-H) and the source level (Figure 3I) even though the subjects performed consistent foot pedaling motion (measured with motion signal monitored by the accelerometer presented in Figure 6). With the modulation of increased UPRF, the MRCP source amplitude was further increased to 2 nA·m. Besides the change of MRCP amplitudes, the emerging timing of negative peak was also postponed from 2 msec in Sham US to 17 msec in UPRF 300Hz condition, and further to 21 msec in the condition of UPRF 3000Hz.

The green patches overlaid on top of the ESI results in Figures 3C, F and I were used to indicate the seed area of the cortical region of interest in our study, i.e. primary leg motor area in left brain hemisphere. We extracted the MRCP source activity from the identified area of 3.4 cm^2^ confined in the reconstructed source area. The resulted MRCP source profiles were generated for subsequent analyses. Figure 4A presents the profiles for comparisons directly among the three ultrasound conditions. In the source domain, typical MRCP features were reconstructed and indicated with horizontal arrows. The readiness potential (RP), also known as Bereitschaftspotential, priors at the onset of movement (from −400 to −24 msec in Sham US and UPRF 300Hz; from from −400 to −20 msec in UPRF 3000Hz) and is involved in the movement preparation (Dremstrup et al., 2013). According to the definition, it is more accurate to denote this MRCP component as “late RP” given its neighboring timing to the movement onset. Immediately following this late RP, the motor potential (MP) reflecting the movement execution was reconstructed within a much short time period of 36-38 msec (from −24 to 12 msec in Sham US; from −24 to 14 msec in UPRF 300Hz; from −20 to 18 msec in UPRF 3000 Hz). Lastly, the MRCP source profile reconstructed the third component, movement-monitoring potential (MMP, from 12 to 178 msec in Sham US; from 14 to 362 msec in UPRF 300Hz; from 18 to 378 msec in UPRF 3000Hz), which controlled the movement performance (Shakeel et al., 2015). Based on the timing of pedaling movement and sonication, the administered tFUS was specifically modulating the MMP. In addition to the significant change on MRCP source profile amplitude (MSPA), the negative source peak was also delayed from 12 msec in Sham US condition to 14 msec and 18 msec in UPRF 300Hz and UPRF 3000Hz conditions, respectively (Figure 4A).

Due to the substantial inter-subject differences during the foot pedaling execution, we firstly conducted the non-parametric paired test for examining the neuromodulatory effects of tFUS on the MRCP in terms of the MSPA. The MSPA is equivalent to the amplitude of MMP as illustrated. When comparing the MSPAs acquired from the Sham US condition (3.13±0.99 nA·m) with those from the UPRF 300Hz condition (6.84±2.39 nA·m), the MSPA was significantly increased (V-statistic = 13, *p* < 0.01). Such a change persisted when the data were also compared against the UPRF 3000Hz condition (15.4±6.89 nA·m, V-statistic = 2, *p* < 0.001). Both tFUS conditions exhibited significantly higher MSPA than the Sham US did.

To further interpret the effect of UPRF in modulating the MRCP, a non-parametric analysis of variance (ANOVA) was performed using Kruskal-Wallis rank sum test (Figure 5A). For this test, the Sham US condition can be considered as a type of tFUS condition with UPRF of 0 Hz. As a result, we found that the UPRF did play a significant role (Kruskal-Wallis chi-squared = 7.24, *p* < 0.05) in changing the MSPA of the human subjects. However, the variance of observations in the condition of UPRF 3000Hz is observed to be larger than the other two conditions. To further probe into a more specific effect of the UPRF, the non-parametric paired Wilcoxon test was further used (Figure 5B). Significantly higher MSPAs (V-statistic = 21, *p* < 0.05) were observed by dosing the higher UPRF, i.e. 3000 Hz.

The MRCP magnitude was correlated with the elbow-flexion movement property, such as the force and its speed (Siemionow et al., 2000). To rule out the confounding factor of induvial subject difference in foot pedaling motion, e.g. the strength and the speed of the leg/foot motion, we compared each individuals’ foot pedaling from the motion signals across all trials as well as the three ultrasound sessions. An example of such comparison is shown in Figure 6. Beyond visual inspection of Figure 6, we further investigated whether the statistical differences among the motions in different ultrasound sessions existed or not by using non-parametric permutation-based statistical tests (5000 randomizations, two-tailed Student’s t-statistic, Bonferroni correction for the multiple comparisons) for all the 1-second trials. No significant difference (*p* > 0.9) was found among different sessions. Similarly, we tested the EMG signal profiles of all valid trials, and still no significant difference was found among those three sessions. With such evidence, one can attribute the observed effect on the MRCPs (Figures 3-5) to the administered tFUS.

## 4. Discussions

In the present study, we introduced the MRCP induced by the real foot pedaling movement as a metric to evaluate the tFUS neuromodulation effect on the human motor cortex. The results demonstrated that tFUS is able to modulate the MRCP in an excitatory way, i.e. increasing the amplitude of MRCP both at the EEG sensor level and at the source domain. One possible mechanism to explain such an enhancement by the tFUS is that the focused acoustic energy may increase excitability of the targeted brain circuit (Gibson et al., 2018; Wang et al., 2019) for a short period. To pursue a high specificity of MRCP readout, we extracted MRCP source activity from individual anatomical brain model, specifically from the primary leg motor area by EEG source imaging. Thus, we are able to reject most of artifacts presented in the electrophysiological recordings with concurrent movement tasks, e.g. unrelated body movements, eye blinks, and event-related sound. Furthermore, we also found that the UPRF of tFUS stimulation plays a positive role in modulation of the MRCP. The excitatory neuromodulation is evidenced in the human primary motor cortex by increasing the UPRF, which is in line with the prediction from computer simulations (Plaksin et al., 2016) and our experimental observations from intracranial recordings at neuronal level in animal models (Yu et al., 2019).

In our experimental paradigm, the tFUS transmission was triggered by the onset of motion signal reflected in the MP phase. Therefore, the 500-msec tFUS was majorly modulating the MMP component of a typical MRCP complex. However, the delayed MMP due to the increased UPRF, i.e. the postponed negative peaks in both sensor (Figures 3A, D and G) and source (Figure 4A) measurements, needs to be further investigated. As illustrated in Figure 4A, the MRCP was initiated during the movement planning phase, i.e. the RP, prior to the actual physical movement. In fact, the RP is raised as early as 1.5 sec before the onset of movement (Shakeel et al., 2015), which may be useful to provide much earlier trigger signal for tFUS neuromodulation, thus assisting the movement planning and preparation. Since the MRCP can also be evoked by motor imagination (Shakeel et al., 2015), the demonstrated neuromodulation by tFUS would lead to more applications in motor rehabilitation/prosthesis scenarios, in which tFUS can be triggered by the early-phase MRCP source activity (i.e. early RP source) of imagining a movement.

Given the importance of the MRCP in the scientific investigations on healthy human subjects (Dremstrup et al., 2013) and clinical evaluations on patients diagnosed with functional motor disorders, such as Parkinson’s disease and Amyotrophic Lateral Sclerosis (ALS) (Gu et al., 2009), it would be valuable to have a non-invasive neuroimaging tool, such as ESI, to map and quantify the MRCP source at specific brain circuits with high spatiotemporal resolution, thus informing the non-invasive neuromodulation for guidance and feedback (Yu et al., 2020). Electrophysiological source imaging (ESI) has been pursued to localize and image brain electrical sources from noninvasive scalp recorded EEG/MEG, and demonstrated to provide greatly enhanced spatial resolution than the raw EEG/MEG in many applications (He & Ding, 2013; He et al, 2018; Sohrabour et al, 2020). The EEG-based ESI (He and Ding, 2013; He et al., 2018) has demonstrated its unique and significant role for effectively identifying epileptogenic brain sources (Sohrabpour et al., 2020) and enhancing the subjects’ BCI performances (Edelman et al., 2019). Although fMRI has been harnessed to monitor and assess the neuromodulatory effects of tFUS on animals models (Folloni et al., 2019a; Verhagen et al., 2019; Yang et al., 2018; Yoo et al., 2011) and on humans (Lee et al., 2016b), the method does not directly measure the neural activities and the strong static magnetic field may confoundingly alter the cortical excitability (Nakagawa and Nakazawa, 2018; Oliviero et al., 2011). In addition, besides the substantial concerns of ultrasound device compatibility with the magnetic field, the fMRI imaging pulse sequences may inevitably induce ancillary brain activations, e.g. auditory responses. Hence, a natural setting for monitoring the tFUS modulating human brain with high spatiotemporal resolutions, like the ESI is desirable.

Low intensity tFUS neuromodulation is deemed as generally safe without inducing serious adverse effects on humans upon careful control of the ultrasound intensities (I_SPPA_: 11.56-17.12 W/cm^2^) (Legon et al., 2020). Although our study administered less ultrasound intensities than those applied in (Legon et al., 2020) and even much less than the FDA’s guideline (I_SPPA_ ≤ 190 W/cm^2^) (FDA, 2008), we were also cautious about the tFUS safety on the subjects during the experiment sessions. We surveyed the subjects with a brief questionnaire for report of symptoms, such as headache, neck pain, dental pain, nausea, dizziness, anxiety, abnormal muscle contractions etc., before and after tFUS sessions. No adverse symptoms have been reported by our human subjects. But in 7 out of all 15 human subjects, scalp tingling sensation was reported only during the UPRF 3000Hz session. Such sensation may distract the subject’s attention on foot pedaling task, which might be the reason for the increased variance of MSPAs observed in this UPRF condition (shown as the UPRF 3000Hz group in Figure 5A).

Due to the time limit in our experiment protocol, we were unable to test extensive UPRF conditions. However, as the UPRF is highly correlated with the ultrasound spatial-peak temporal-average intensity (I_SPTA_), by increasing this frequency, the total acoustic energy deposited onto the targeted brain area is correspondingly increased. It is reasonable to infer that the increased ultrasound energy in a high UPRF condition leads to increased excitatory effects at the human primary motor cortex. Future research efforts will adjust UPP for different UPRF levels to maintain the total ultrasound energy as a constant during the fixed sonication period to tease out whether UPRF frequency encoding plays a role in the modulation effects.

Inevitably, we observed inter-subject differences in response to the tFUS neuromodulation (shown in Figure 4B-C and Figure 5B). Such differences may be attributed to many factors. One of the crucial factors is the inter-subject variation in skull morphology and composition. No linear relationship was found either between the skull thickness and the transcranial maximal acoustic pressure or between the skull thickness and the ultrasound full-width at half magnitude volume behind the skull (Mueller et al., 2017). To address this issue, tFUS dose and target planning before the neuromodulation experiment would be required and should be individualized; such a planning may rely on *a priori* computer simulations for predicting acoustic fields within individual skull cavity based on respective skull model established from CT scans (Mueller et al., 2017; Webb et al., 2018).

## Acknowledgements

This work was supported in part by NIH grants EB029354, MH114233, AT009263, EB021027, and NS096761. X.N. was supported in part by Carnegie Mellon Neuroscience Institute Presidential Fellowship and Liang Ji Dian Graduate Fellowship at Carnegie Mellon University.

We thank Dr. Abbas Sohrabpour, Dr. Haiteng Jiang and Daniel Suma for useful discussions, Shahriar Noroozizadeh, Emily Lopez, Xiyuan Jiang and Sandhya Ramachandran for experimental assistance. We would also thank the technical support from the CMU-Pitt Brain Imaging Data Generation & Education (BRIDGE) Center for providing the MRI facilities and services.

## Declaration of Interests

The authors declare no competing interests.

## Data and Software Availability

All data and software are available upon reasonable request to Bin He.

## Notes

### Competing Interest Statement

The authors have declared no competing interest.

